# SYNERGISTIC ANTITUMOR EFFECT OF NAPROXEN AND SORAFENIB IN HEPATOCELLULAR CARCINOMA

**DOI:** 10.1101/2024.02.01.578341

**Authors:** Etkin Akar, Seyma Unsal Beyge, Deniz Cansen Kahraman

## Abstract

Hepatocellular carcinoma (HCC) is one of the most prevalent and deadliest cancer types in the world. Due to the inevitable development of resistance to classical chemotherapy and current targeted therapies, combination therapy became an important strategy. Recently drug repurposing studies have emphasized the potential of non-steroidal anti-inflammatory drugs (NSAIDs) in exhibiting anti-tumor activities in different cancer types, including HCC. In this study we explored the synergistic bioactivity of naproxen in combination with sorafenib in HCC cell lines. Network reconstruction analysis, performed to identify key factors mediating the synergistic therapeutic effects of sorafenib and naproxen revealed apoptosis pathways enriched in both sorafenib and naproxen networks. CAPN2 emerged as the common gene shared between the two networks. *In vitro* studies on Huh7 and Mahlavu cells revealed that induction of apoptosis was associated mainly with the PI3K/AKT pathway inhibition. Furthermore, combined treatment of naproxen and sorafenib led to the downregulation of CAPN2 expression, suggesting a potential link to apoptosis and PI3K/AKT pathway regulation, which needs further investigation. In conclusion, combining sorafenib with naproxen presents a promising strategy with potential synergistic therapeutic benefits in HCC.

## INTRODUCTION

Hepatocellular carcinoma (HCC) is the most common form (90%) of primary liver cancers and is the second leading cause of cancer-related death worldwide. Approximately, 850,000 people are diagnosed with liver cancer annually. Chronic infection with Hepatitis B (HBV) and C virus (HCV) infection, alcohol intake, aflatoxin B1, and nonalcoholic steatohepatitis (NASH) are potential risk factors for HCC (1). Patients with early or intermediate-stage HCC can be cured by hepatectomy, liver transplantation, and local ablation. However, advanced-stage HCC patients can only benefit from systemic therapy. As targeted therapy for HCC has been extensively investigated, accumulating evidence has highlighted the significant synergistic anti-tumor effect of multi-target combination therapy. Sorafenib, a multi-target kinase inhibitor possessing anti-angiogenic and anti-proliferative effects, has been shown to increase overall survival (OS) in patients with advanced HCC from 8 to 11 months and was the sole systemic therapeutic agent approved for HCC between 2007 and 2016 (2). In 2018, lenvatinib was approved as the second first-line treatment for patients with advanced HCC (3), while regorafenib received FDA approval in 2017 as a second-line treatment for individuals with unresectable HCC (4). Additionally, in 2019, cabozantinib was utilized as a second-line treatment for those previously treated with sorafenib (5). Whilst systemic therapy may extend the median survival period of patients with advanced HCC, the emergence of tumor cell resistance to kinase inhibitors often results in treatment failure. This has emerged as a significant impediment to the successful clinical management of advanced HCC patients. Hence, it is imperative to identify dependable drug candidates that may offer new therapeutic avenues for patients with advanced HCC.

Nonsteroidal anti-inflammatory drugs (NSAIDs) have been for many used in the management of acute and chronic conditions characterized by pain and inflammation. Lately, clinical investigations have underscored the potential of NSAIDs in the treatment of cancer, with evidence suggesting that they are efficacious in reducing the incidence and mortality of several types of cancers, including HCC (6–8). Numerous preclinical and clinical investigations have reported that co-administration of chemotherapeutic agents and anti-inflammatory drugs can significantly enhance patient prognosis. While monotherapy may not completely eradicate cancer, anti-inflammatory agents may serve as valuable adjuvants to conventional therapeutic approaches (9). Given that over 90% of liver cancer cases have been linked to chronic inflammation, nonsteroidal anti-inflammatory drugs (NSAIDs) may represent a promising therapeutic option for HCC. Over the past decade, the anticancer and anti-metastatic effects of aspirin, a specific type of nonsteroidal anti-inflammatory drug (NSAID), have been extensively documented in various cancers, including HCC (10, 11). One study revealed that aspirin can trigger apoptosis via mitochondrial dysfunction in HepG2 cells by inducing metabolic and oxidative stress (12). Another investigation demonstrated that aspirin could alleviate sorafenib-induced pro-metastasis effects by enhancing the expression of the tumor suppressor HTATIP2 in nude mice xenografts (13). In addition, one of our recent studies presented the *in vitro* and *in vivo* anticancer properties of a triazolothiadiazine derivative NSAID against HCC cells and demonstrated the anti-stemness effect of this derivative in combination with sorafenib (14). Thus, it is vital to investigate the benefits of combining multikinase inhibitors with NSAIDs to improve the clinical outcome of patients with advanced HCC.

Protein-protein and drug-protein interactions play a crucial role in understanding which pathways are activated or inactivated by the administration of drug(s) within the cell. As drugs perturb multiple signaling pathways along with their targets, inference of modulated pathways helps us to understand the underlying disease mechanisms and to find out similarities and/or differences between the effects of different drugs. With the aid of network-based analysis of omics data, it is possible to discover hidden targets that are commonly affected by drugs having different mechanisms of action. In combinatorial treatment strategies, it is suggested that drug-modulated networks should have commonalities on the altered proteins, while they should also affect different molecular pathways to be more effective against the disease (15). Thus, it is essential to investigate the network-level alterations of drugs to understand their potential synergistic effects.

In this study, we demonstrated the synergistic effects of naproxen and sorafenib for the first time on HCC cell lines and investigated the possible underlying mechanisms through network-based analysis and *in vitro* assays.

## MATERIALS AND METHODS

### Cell Culture

Mahlavu (16) and Huh7 (JCRB0403) cell lines were grown in Dulbecco’s Modified Eagle Medium (DMEM, cat no: BI01-050-1A) supplied with %10 FBS (cat no: 10270), Penicillin/streptomycin (cat no: 15140-122), L-glutamine (cat no: 25030), from Thermo Fisher Scientific, 37 °C under 6% CO2. The cell lines utilized in this research have undergone authentication through short tandem repeat (STR) analysis. Routine examinations for mycoplasma contamination were conducted using a mycoplasma detection kit (MycoAlert™, Lonza, cat. no. LT07-318). The cells were passaged no more than 8-10 times (twice a week) during the experiments.

### Sulforhodamine B (SRB) assay

Huh7 (2500 cells/well) and Mahlavu cells (1500 cells/well) were seeded in 96-well plates in 150 µl complete media for 24h and treated with 10 µM to 0.62 µM of sorafenib, lenvatinib, regorofenib and/or 400, 200 and 100 µM naproxen, flurbiprofen, ibuprofen and aspirin. After 72h of treatment, cells were washed with 1X PBS and fixed with ice-cold 10% trichloroacetic acid (Sigma Aldrich; Merck cat. no: 27242) at dark, +4 C, for 1h, then washed with ddH_2_O. Air-dried plates were stained with SRB dye dissolved in %1 acetic acid (Sigma-Aldrich, cat. no: 27725) for 15 minutes and then rinsed with %1 acetic acid. A 10 mM Tris-base solution is applied to air-dried plates for 15 minutes, with gentle shaking to solubilize the protein. Subsequently, the plates are read at 515 nm using a BMG Labtech SuperStar Nano plate reader.

### Synergyfinder Tool

*Synergyfinder* web-based tool (17) was used to quantify the degree of synergism between sorafenib and naproxen. Percent inhibition values for each treatment condition were calculated as follows:

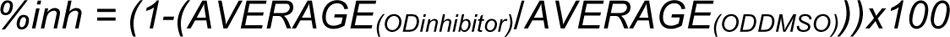

ZIP (Zero interaction potency model), which characterizes drug interaction connections by examining the variations in potency (effect at a specific dose level) within the dose–response curves of individual drugs and their combinations, is used to assess the synergy score for each combination.

### Real-time cell growth analysis (RT-CES)

Mahlavu and Huh7 cells were plated into E-96 plates (Agilent, cat. no:5232368001), 1500-2500 cells/wells, respectively. The next day, cells were treated with the selected concentration of the drugs. Fast drug responses were recorded every 10 minutes, then long-term drug responses were recorded every 30 minutes through (RT-CES) (ACEA Biosciences; Agilent Technologies). Cell Index values (CI) were employed to construct the real-time proliferation curve for each condition by utilizing time-zero normalized CI values.

### Pathway analysis and network reconstruction analysis

Transcriptomic data of naproxen was collected from Connectivity Map (CMap) LINCS 2020-level5 data (18), and sorafenib data was obtained from E-MTAB-7847 RNAseq data. Differentially expressed genes from common gene sets were combined with corresponding drug targets collected from Cmap Drug Repurposing Tool (19). Then, each gene list was used to reconstruct drug-disease networks using Omics Integrator-Forest tool (20). Further pathway enrichment analysis was performed to analyze commonalities in drug-targeted networks.

### Apoptosis assay by Annexin-V/PI staining

Mahlavu and Huh7 cells were plated in 6-well plates (30-60×10^3^ cells/wells), respectively. The next day, cells were treated with selected concentrations of sorafenib and naproxen. After 72h of incubation, cells were collected via trypsinization,centrifuged at 1800 rpm for 5 minutes, and rinsed twice with 1x PBS. For Annexin V/PI staining, BD Pharmingen™ FITC Annexin V Apoptosis Detection Kit I (BD, cat. no: 556547) was used as recommended by the manufacturer. Briefly, cell pellets were resuspended with 1X assay buffer (100 μl/10^6^ cells) supplemented with 1% of FBS. Then, cells were stained with 1 µl of PI and 1 µl of Annexin-V dye, followed by 15 minutes of incubation at RT. 400 µl of 1xPBS was added to each sample before analysis using Novocyte flow cytometry (ACEA Biosciences; Agilent Technologies). Apoptotic cells were determined by NovoExpress software (ACEA Biosciences; Agilent Technologies).

### Immunofluorescence staining

Mahlavu and Huh7 cells were seeded onto cover slides placed in 6-well plates (30-60×10^3^ cells/wells). The next day, cells were treated with selected concentrations of drugs. At the end of each time point, cells were fixed with %100 cold methanol for 10 minutes, then washed with 1xPBS multiple times. Cell nuclei were stained with 1 μg/ml Hoechst 33258 dye for 5 minutes at RT. Then, the excess dye was removed by 10 minutes of incubation with ddH_2_O at RT. Apoptotic nuclei were identified using a blue filter (340–380 nm) on a fluorescence microscope (Nikon Eclipse 50i).

### Western Blot analysis

Mahlavu cells were seeded in 15 mm dishes, 60×10^5^ cells/dishes, respectively. The next day, cells were treated with the synergistic concentrations of sorafenib and naproxen (Sor: 5 μM, Nap: 400 μM, or Nap+Sor (5 μM + 400 μM). After 24h, cells were harvested by scraping and washed with 1xPBS. Cells were lysed with NP-40 lysis buffer by centrifugation at 13000 rpm for 30 minutes. Protein concentration was determined by BCA assay (Sigma Aldrich, cat no: B9643). 30-40 μg protein was loaded in Mini-PROTEAN® Tetra Cell Systems and TGX™ precast gels. Samples were run in Tris/Glycine/SDS (TGS) Running Buffer, (Bio-Rad, 161-0772). Trans-Blot® Turbo Transfer System (Bio-Rad) was used to transfer proteins from gel to the LF-PVDF membrane (Bio-Rad, cat. no: 1704274). For immunoblotting, the blots were incubated with primary antibodies against PARP (Cell Signaling Technology, Danvers, MA, USA, cat. no: 9532S, 1:1000). Caspase-9 (Santa Cruz Biotechnology, cat. no: sc-22182, 1:100), Caspase-8 (Cell Signaling Technology, Danvers, MA, USA, cat. no: 9746S, 1:100), Bcl-xS/L (Santa Cruz Biotechnology, cat. no: sc-634, 1:100), Cytochrome-c (Santa Cruz Biotechnology, cat. no: sc-7159, 1:100), mTOR (Cell Signaling Technology, cat no. 2983S, 1:250), phospho-mTOR (Ser2448) (Cell Signaling Technology, cat. no: 2971S, 1:250), Akt (Cell Signaling Technology, cat. no: 9272S, 1:500), phospho-Akt (Ser473) (Cell Signaling Technology, cat. no: 9271S, 1:500), Calnexin (Cell Signaling Technology, cat. no: 2679S, 1:2000), Actin (Cell Signaling Technology, cat. no: 4967S, 1:2000). After the primary antibody incubation, membranes were washed with 1X TBS-T four times and incubated with secondary antibody including 680RD anti-rabbit (Li-cor, cat. no: 926-32211 1:10.000), 680RD anti-mouse (Licor, cat. no: 926-32210, 1:10.000), 800CW anti-goat (Licor, cat. no: 925-32214, 1:10.000) for 1h at RT in dark. After washing with 1X TBS-T three times, proteins were visualized in the LI-COR Odyssey CLx imaging system and analyzed using Image Studio Lite Ver 5.2.

### Quantitative RT-PCR (qRT-PCR)

Total RNA was isolated from cells using the RNeasy RNA-purification kit (Qiagen, cat. no:74106), and cDNA was synthesized by using the RevertAid First Strand cDNA synthesis kit (Fermantas, cat. no: K1622) The primer sequences specific for CAPN2 and the internal control gene RPL-19 were designed and used for the qPCR analysis. CAPN2 forward primer: 5’TGCTCCATCGACATCACCAG 3’, CAPN2 reverse primer: 5’GTCTGGTCAGCCTTTCCCTC 3’, RPL19 forward primer 5’GCTCTTTCCTTTCGCTGCTG 3’, RPL19 reverse primer 5’GGATCTGCTGACGCGAGTTG 3’. Light Cycler® 96 Real-Time PCR System (Roche) with LightCycler® 480 SYBR Green I Master (Roche, cat. no:04 707 516 001) was used to perform quantitative PCR under following conditions: initial denaturation for 10 min at 95 °C, 45 cycles amplification for 10 sec at 95°C followed by 10 sec at 61°C and 10 sec at 72°C, and melting for 10 sec at 95°C followed by 65°C for 60 sec and 1 sec at 97°C. The experimental CT is calculated compared to internal control RPL19 products. The ΔΔCt was used to determine relative expression of the target genes compared to the control group.

### Statistical analysis

All the experiments were carried out with n≥3 biological replicates. One-way ANOVA and Student’s t-test were applied to determine significance. * p<0.05, ** p<0.01, *** p<0.001

## RESULTS

### Naproxen and Sorafenib exhibit synergistic interaction in HCC cell lines

First, we examined the anti-cancer activity of FDA-approved NSAIDs, including aspirin, ibuprofen, flurbiprofen, and naproxen, in combination with the HCC drugs, including sorafenib, regorafenib, and lenvatinib by SRB assay. Cytotoxicity served as the basis for evaluating the interaction pattern of each combination using the *Synergyfinder* tool. Among all combinations (**Table S1**), naproxen and sorafenib demonstrated a synergistic interaction in HCC cell lines, with scores of 16.142 and 16.482 for Mahlavu and Huh7 cells, respectively (**Fig. 1A**). The xCELLigence system was utilized to analyze the time- and dose-dependent growth pattern of Mahlavu and Huh7 cells treated with synergistic concentrations of sorafenib and naproxen which were determined based on the previously defined synergistic area. The sorafenib-naproxen combination significantly reduced the growth of HCC cell lines in a dose-dependent manner compared to sorafenib alone. (**Fig. 1B**). The most potent synergistic concentrations for the naproxen-sorafenib combination were determined as 2.5 μM sorafenib + 400 μM naproxen for Huh7, and 5 μM sorafenib + 400 μM naproxen for Mahlavu cells. Moreover, to assess the morphological changes induced by this combination, cells were treated with these synergistic concentrations and visualized under light microscopy. The number of detached cells significantly increased, resulting in reduced overall viability (**Fig. S1**).

**Figure 1.**
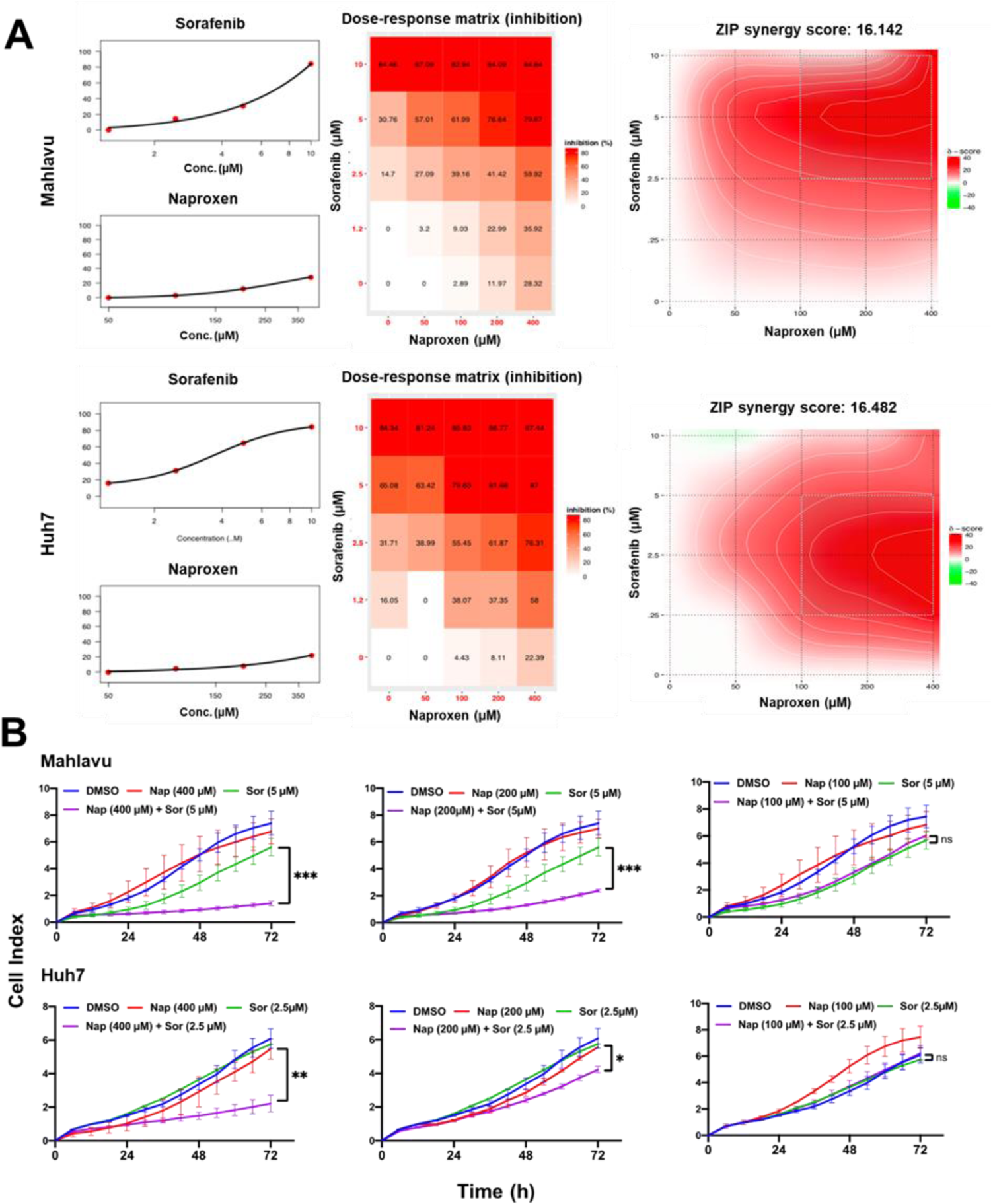
Naproxen and sorafenib are synergistically interacting and induce dose-dependent cell growth inhibition on HCC lines. (**A**) The cytotoxicity of each drug and their combination were evaluated by NCI-SRB assay. The dose-response curves and matrices, as well as the heatmaps were generated after 72h of treatment. Synergy scores were calculated by the zero-interaction potency model (ZIP) by *Synergyfinder*. The most synergistic areas were shown in dark red (**B**). The real-time cell growth curves of the selected concentrations of the naproxen-sorafenib combination, and their control DMSO were generated by the RT-CES system. Cell index (CI) was recorded every 30 minutes. All treatments were done in triplicates.

### Network analysis revealed enrichment of apoptotic pathways in HCC cells treated with the combination of sorafenib and naproxen

To reveal pathways implicated in sorafenib and naproxen treatment in HCC, we analyzed sorafenib and naproxen transcriptomic data sets, which are publicly available in OMICSDI-EMTAB-7847 and CMAP databases, respectively. Network analysis was performed to investigate the possible hidden mechanisms of the combination. It allows the integration of not only known targets of the drugs but also hidden pathways modulated by the treatments. The pipeline of the analysis is shown in (**Fig. 2A**). Although sorafenib and naproxen mostly modulate different pathways and only a single gene (CAPN2) was found common in both networks, apoptotic pathways were enriched in Huh7 cells co-treated with sorafenib and naproxen. (**Fig. 2B and 2C**).

**Figure 2.**
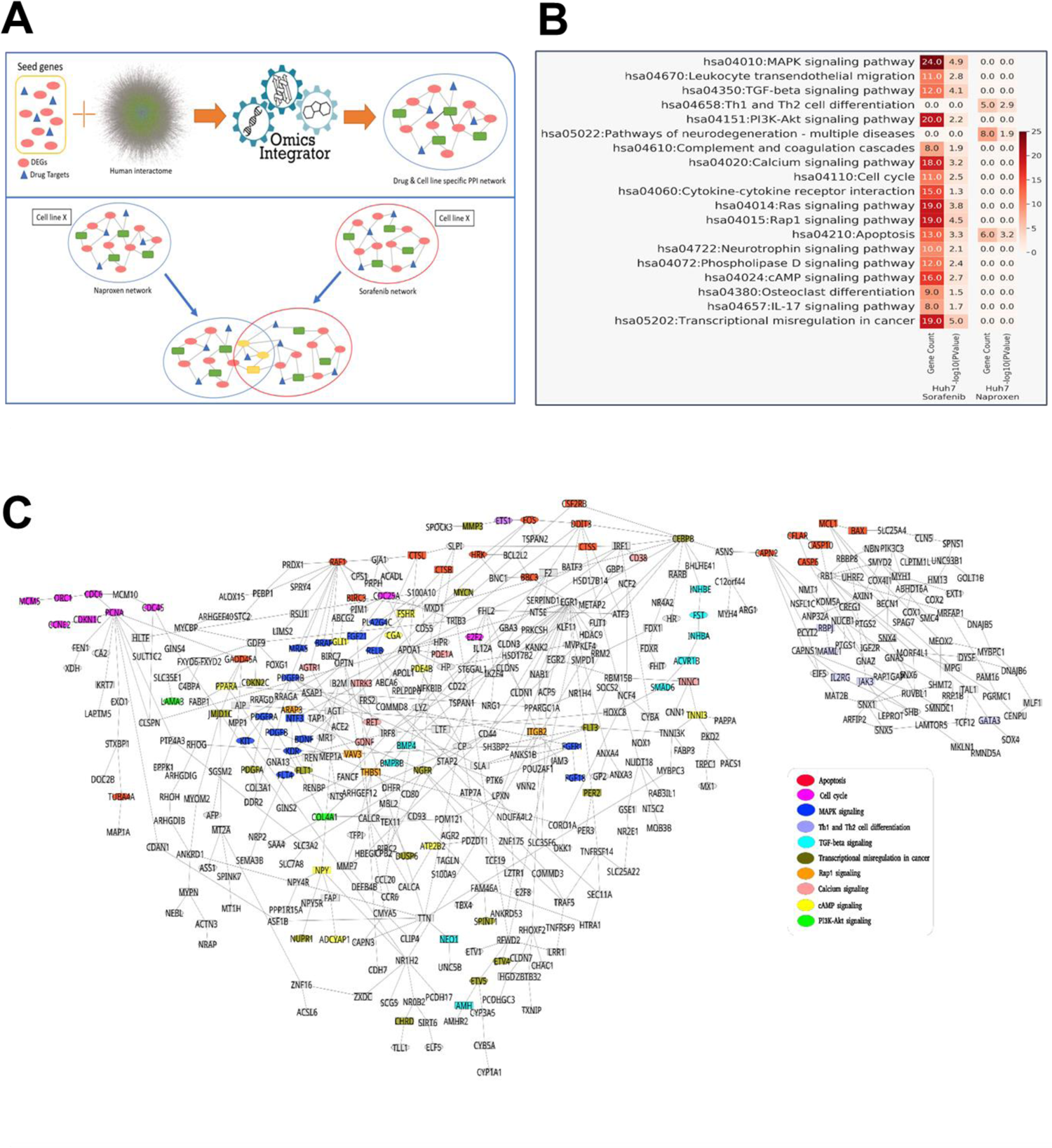
Apoptotic pathways are enriched in both sorafenib and naproxen networks in Huh7 cells. (**A**) Illustration of the transcriptomics dataset pipeline for Huh7 cells treated with either sorafenib or naproxen. (**B**) Results of pathway enrichment for the drugs. (**C**) Network analysis of sorafenib and naproxen, with node colors indicating the enriched pathways where the gene is active. It is essential to note that a single gene may appear in multiple pathways, but the prioritization for coloring is determined by the significance of pathway enrichments.

### Sorafenib and naproxen combination induces apoptosis in HCC cell lines

Since apoptotic pathways were shown to be implicated in both networks, we focused on apoptosis in HCC cell lines. Flow cytometric analysis of Annexin-V/PI-stained cells revealed that the apoptotic cell population significantly increased in the sorafenib-naproxen combination compared to the control and single-drug treated groups (**Fig. 3A**). Moreover, Hoechst staining and immunofluorescence imaging of HCC cells have shown condensed chromatin structures and nuclear blebbing were increased upon treatment with the sorafenib-naproxen combination compared to the other groups (**Fig. 3B**). Lastly, expression of pro-apoptotic proteins such as cleaved-PARP, cleaved-caspase-9, cleaved-caspase-8, and BCL-x S/L increased significantly upon sorafenib-naproxen combination compared to sorafenib treatment (**Fig. 3C**).

**Figure 3.**
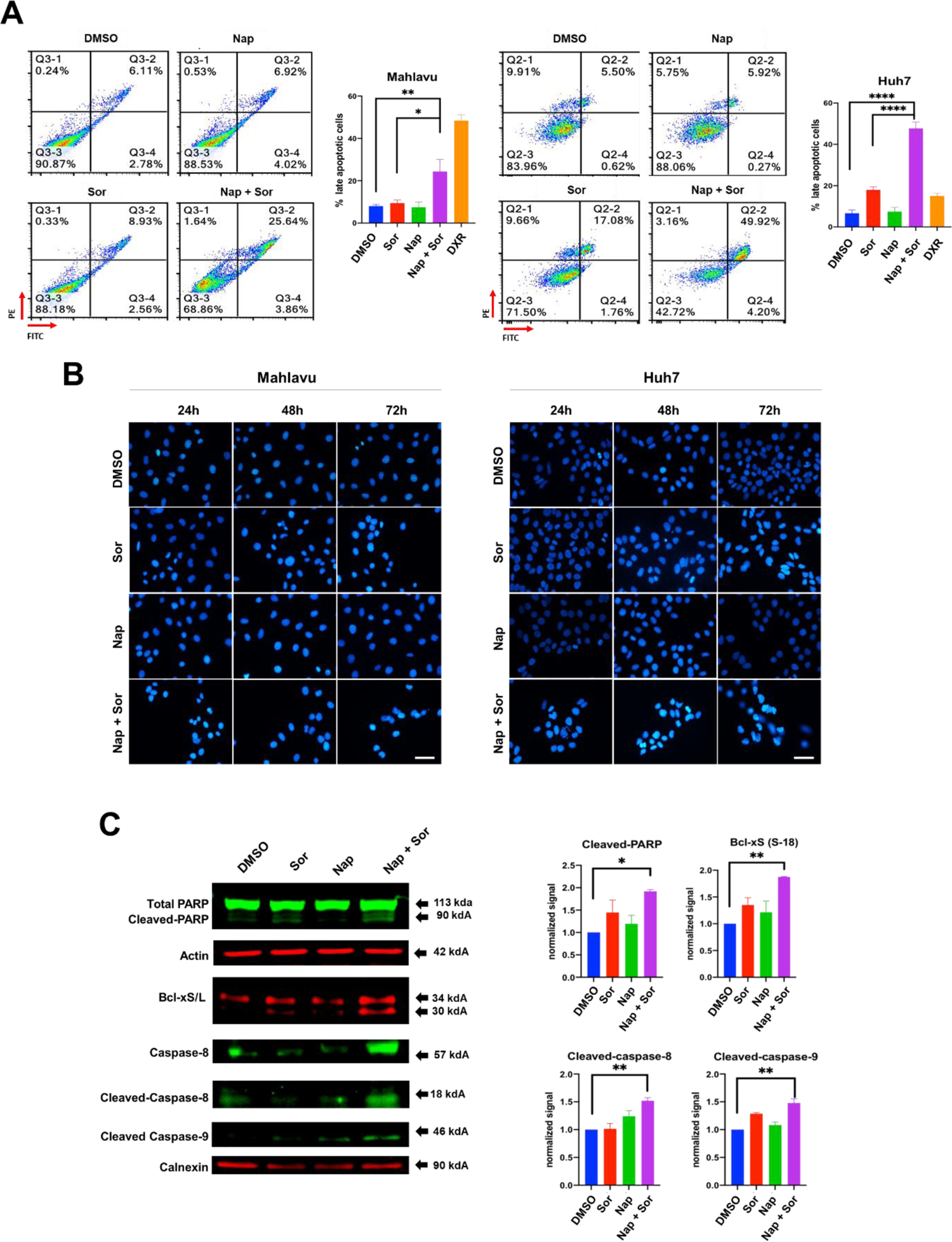
Sorafenib and naproxen combination induce apoptosis in HCC cell lines. (**A**) HCC cells were treated with Sor: 5 μM for Mahlavu, 2.5 μM for Huh7, Nap: 400 μM, or Nap+Sor (5 μM + 400 μM for Mahlavu, and 2.5 μM + 400 μM for Huh7) for 72h and stained with BD Pharmingen™ FITC Annexin V Apoptosis Detection Kit to be analyzed using flow cytometry. Bar graphs indicate % late apoptotic cells in different treatment groups. DXR: Doxorubicin, used as a positive control (0.1 μM) for apoptosis induction. (**B**) Fluorescence microscopy images of Mahlavu and Huh7 cells treated with the same concentrations as in part A for 24h, 48h, or 72h, followed by staining with Hoechst 33258. Condensed nuclei and nuclear blebbing are observed in a light-blue color. (scale bar: 50 µm) (**C**) Western blot analysis of apoptotic proteins in Mahlavu cells. Actin and calnexin were used as loading controls. Bar graphs indicate relative protein expression for each protein in different treatment groups. Sor: Sorafenib, Nap: Naproxen.

### *CAPN2* is downregulated upon treatment with sorafenib and naproxen combination

*CAPN2* is the only gene common in both sorafenib and naproxen networks (**Fig. 2C**), which may be one of the key elements in the synergistic activity of the sorafenib-naproxen combination. *CAPN2* is reported to be upregulated in various cancer types, including HCC, and it is known to promote invasion and metastasis (21), (22). In the analysis of overall survival among cancer patients, high *CAPN2* expression was associated with a significant decrease in survival rates (**Fig. 4A**). Furthermore, a TNM plot revealed a noteworthy increase in *CAPN2* expression levels in both primary liver tumors and metastatic liver tumors (**Fig. 4B**). These findings underscore the prognostic significance of elevated *CAPN2* levels in cancer patients and highlight its potential role in liver cancer progression and metastasis. Therefore, we checked the mRNA levels of *CAPN2* in HCC cell lines by quantitative PCR (qPCR). Strikingly, *CAPN2* expression was upregulated upon sorafenib treatment. Notably, the co-treatment of sorafenib and naproxen resulted in a significant and dramatic reduction in *CAPN2* expression in both HCC cell lines (**Fig. 4C**). This observation highlights the potential therapeutic impact of the sorafenib and naproxen combination on the regulation of *CAPN2* expression, which merits further exploration in the context of HCC.

**Figure 4.**
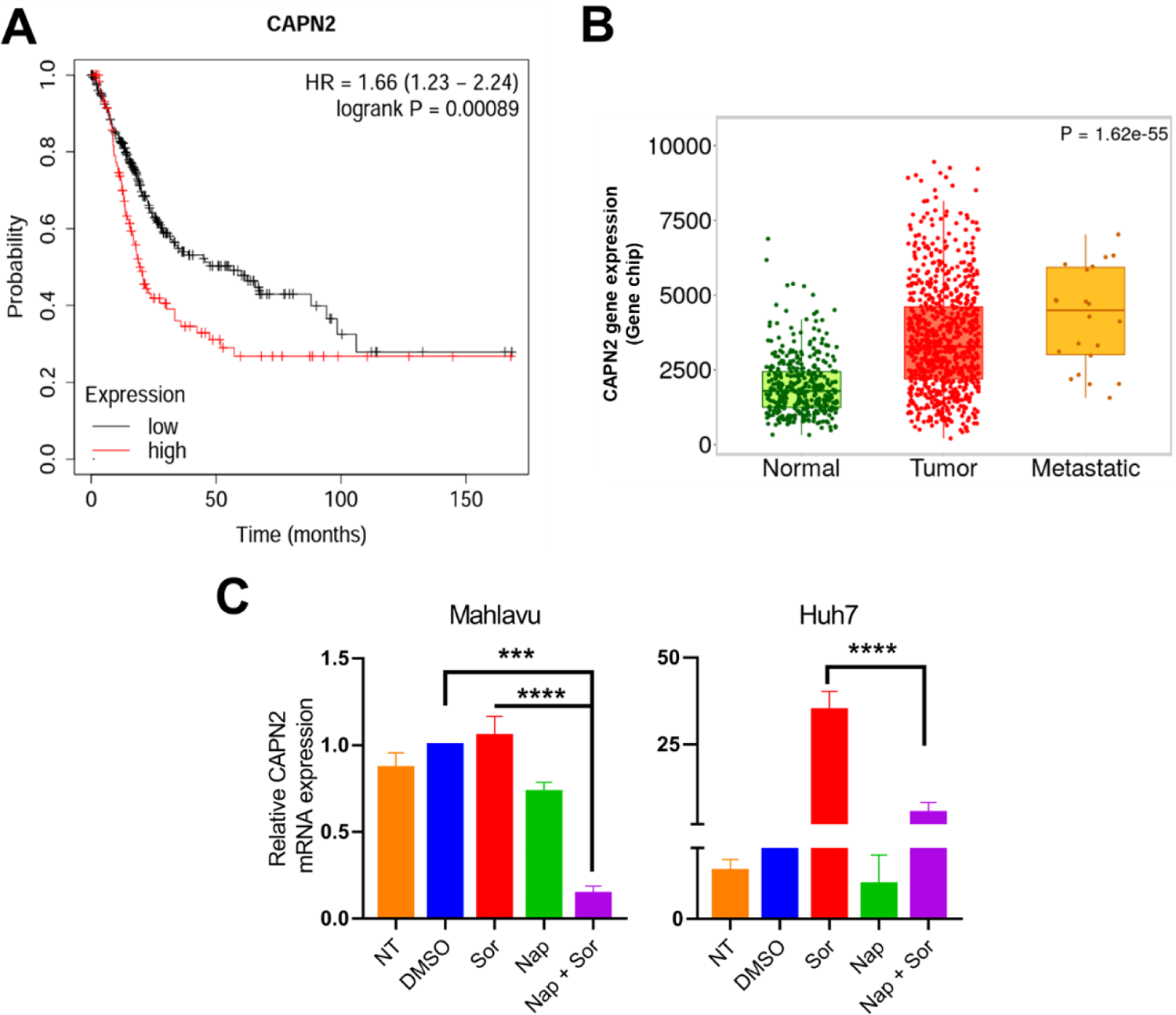
*CAPN2* expression in HCC is downregulated by combinatorial treatment of sorafenib and naproxen. (**A**) Kaplan Meier survival plot generated using *Kaplan-Meier plotter* (23), showing a significant decrease in overall survival among cancer patients with high *CAPN2* expression (**B**) TNM plot generated using the *TNM plot tool* (24), showing a notable increase in *CAPN2* expression levels in both primary liver tumors and metastatic liver tumors. (**C**) Bar graphs indicate relative mRNA expression of *CAPN2* in Mahlavu and Huh7 cells treated with Sor: 5 μM for Mahlavu, 2.5 μM for Huh7, Nap: 400 μM, or Nap+Sor (5 μM + 400 μM for Mahlavu, and 2.5 μM + 400 μM for Huh7) for 48h (NT: no treatment, Sor: Sorafenib, Nap: Naproxen)

### Sorafenib and naproxen combination decreases phosphorylation of mTOR and AKT proteins

The PI3K/AKT pathway, known for its hyperactivity in various cancer types, including HCC, plays a pivotal role in cell survival and apoptosis regulation (25). To elucidate the impact of sorafenib and naproxen combination on this critical signaling cascade, we conducted western blot analysis using Mahlavu cells, given the well-established hyperactivity of the PI3K/AKT pathway in these cells (26). Notably, the co-treatment of sorafenib and naproxen led to a significant reduction in the phosphorylation levels of AKT and mTOR, key effectors of the PI3K/AKT pathway (**Fig. 5**).

**Figure 5.**
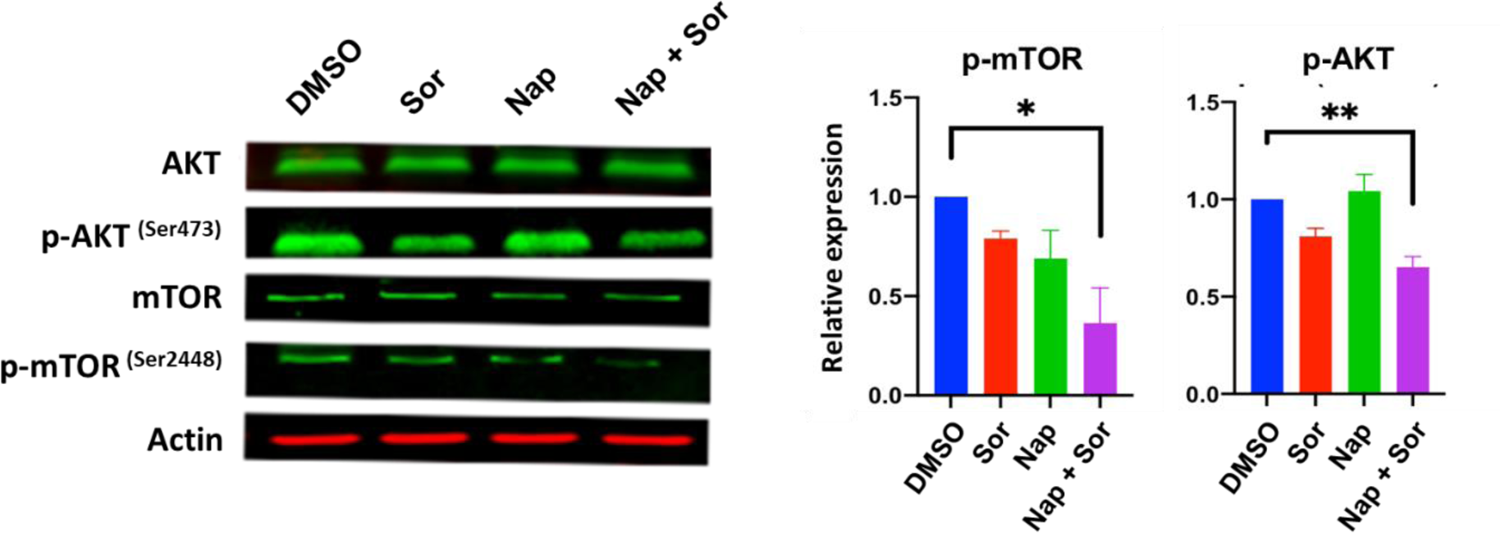
PI3K-AKT pathway is down-regulated in HCC by sorafenib and naproxen combination. Western Blot analysis of total and phosphorylated AKT (Ser473) and mTOR (Ser2448) proteins in Mahlavu cells treated with Sor: 5 μM, Nap: 400 μM, or Nap+Sor (5 μM + 400 μM) for 24 h (NT: no treatment, Sor: Sorafenib, Nap: Naproxen). Actin was used as a loading control.

## DISCUSSION

This study reveals a novel synergistic interaction between the FDA-approved drugs sorafenib and naproxen in HCC. Notably, this marks the first instance of establishing a synergistic relationship between these drugs for HCC treatment. Prior studies on other cancer types, such as those by Kim et al. (27) and Xia et al. (28), utilized higher concentrations of NSAIDs, specifically 2 mM or 4 mM. In contrast to these existing findings, our study demonstrates synergistic effects at lower concentrations on HCC cell lines, which could potentially minimize adverse effects and enhance the safety profile of the treatment.

To unravel the signaling pathways that could underpin this synergistic effect, we employed transcriptomic datasets of sorafenib and naproxen in HCC cell lines for network reconstruction analysis. This approach facilitated the discovery of hidden targets of the drugs, extending beyond their primary targets, and illuminated the signaling pathways modulated within this interactive synergy. Our results revealed the enrichment of apoptotic pathways in both sorafenib and naproxen networks, which was further validated *in vitro* on HCC cell lines. Pro-apoptotic proteins were significantly upregulated in HCC cells upon treatment with both drugs. Moreover, the downregulation of PI3K/AKT pathway proteins such as phospho-AKT and phospho-mTOR suggested a plausible mechanistic link between the drug combination and the suppression of pro-survival signals.

Moreover, *CAPN2* gene, which was found to be the only gene mediated by both sorafenib and naproxen network, was downregulated when two drugs were given in combination to HCC cell lines. Research has demonstrated that elevated CAPN2 expression is linked not only to cancer proliferation and metastasis, as indicated by studies such as those by Miao et al. (21) and Peng et al., (29), but is also correlated with reduced survival rates in cancer patients, including those with HCC. Moreover, CAPN2 was recognized as the central point within a protein-regulatory network associated with aggressive phenotypes in HCC (30). Our qPCR analysis showed that Huh7 cells exhibit notably low levels of CAPN2 expression, an observation consistent with data from the Harmonizome database (31). In contrast, Mahlavu, characterized as one of the most aggressive HCC cell types, demonstrates a markedly higher baseline expression of CAPN2. Furthermore, a consistent pattern is observed across both cell lines: sorafenib induces an increase, naproxen leads to a decrease, and the combined administration of the drugs significantly reduces CAPN2 expression. This suggests its involvement in the molecular pathways influenced by this synergistic drug combination. CAPN2 silencing has been shown to promote apoptosis in non-small cell lung cancer (NSCLC) (32) and many studies address CAPN2 down-regulation associated with decreased activity of PI3K/AKT pathway in other cancer types (21, 33, 34). Our results also pointed out that apoptosis induction may be further associated with the downregulation of the CAPN2 gene and the decreasing activity of the PI3K-AKT pathway in HCC (**Fig. 6**). Yet, future research could explore the functional role of CAPN2, providing a deeper understanding of its mechanisms and its link to the observed synergistic effect. Additionally, to validate the translational potential of our *in vitro* findings, it is imperative to conduct *in vivo* studies that assess the synergistic effect of sorafenib and naproxen. This step will provide crucial insights into the feasibility and efficacy of the drug combination in a more complex physiological context, ultimately advancing its potential as a therapeutic strategy for HCC.

**Figure 6.**
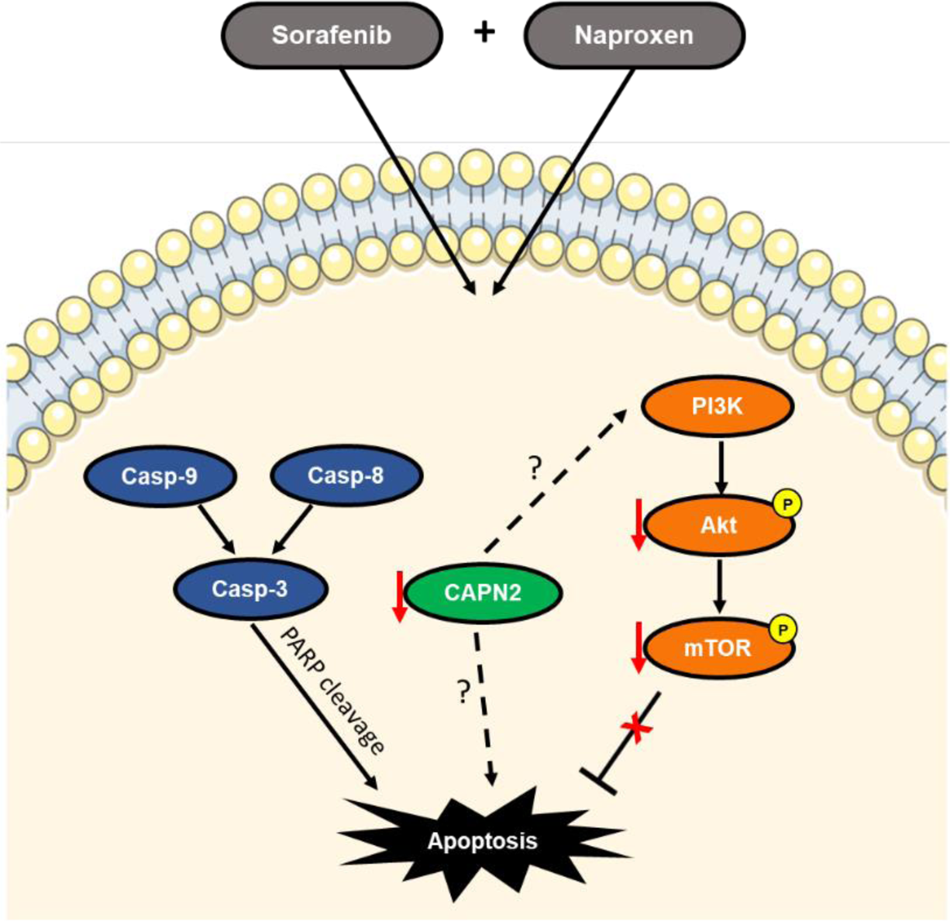
Sorafenib and naproxen co-treatment induces activation of pro-apoptotic proteins and inhibition of PI3K/AKT pathway proteins in HCC. The underlying mechanisms involved in this synergistic effect are activation of pro-apoptotic proteins and inhibition of PI3K/AKT pathway proteins. Downregulation of CAPN2 might serve as an important link between these mechanisms.

Overall, our study demonstrates that the combination of sorafenib and naproxen induces apoptosis in HCC cell lines through the inhibition of the PI3K/AKT pathway and activation of pro-apoptotic proteins. The identification of CAPN2 as a common gene in the sorafenib and naproxen networks suggests its potential role as a crucial element in this synergistic interaction. These findings not only contribute to our understanding of the molecular mechanisms underlying the anticancer effects of this drug combination but also suggest potential therapeutic targets for HCC.

## Acknowledgments

We would like to express our sincere gratitude to Dr. Nurcan Tuncbag for her invaluable guidance and insightful comments throughout the development of this manuscript. We would also like to thank Dr. Esra Nalbat for her proofreading and editing contributions. This work was supported by The Turkish Ministry of Development project (#KanSil 2016K121540).

## Author contributions

DCK conceived and supervised the study. EA performed the *in vitro* experiments. SUB exhibited the *in silico* network analysis. DCK, EA, and SUB wrote the initial draft of the manuscript. All authors contributed to the interpretation of results, critically reviewed, edited the manuscript, and approved the final version for submission.

## Notes

### Competing Interest Statement

The authors have declared no competing interest.

## References

1. Llovet JM, Kelley RK, Villanueva A, Singal AG, Pikarsky E, Roayaie S, et al. Hepatocellular carcinoma. Nat Rev Dis Primers. 2021;7(1):6.

2. Llovet JM, Ricci S, Mazzaferro V, Hilgard P, Gane E, Blanc JF, et al. Sorafenib in advanced hepatocellular carcinoma. N Engl J Med. 2008;359(4):378–90.

3. Kudo M, Finn RS, Qin S, Han KH, Ikeda K, Piscaglia F, et al. Lenvatinib versus sorafenib in first-line treatment of patients with unresectable hepatocellular carcinoma: a randomised phase 3 non-inferiority trial. Lancet. 2018;391(10126):1163–73.

4. Bruix J, Qin S, Merle P, Granito A, Huang YH, Bodoky G, et al. Regorafenib for patients with hepatocellular carcinoma who progressed on sorafenib treatment (RESORCE): a randomised, double-blind, placebo-controlled, phase 3 trial. Lancet. 2017;389(10064):56–66.

5. Abou-Alfa GK, Meyer T, Cheng AL, El-Khoueiry AB, Rimassa L, Ryoo BY, et al. Cabozantinib in Patients with Advanced and Progressing Hepatocellular Carcinoma. N Engl J Med. 2018;379(1):54–63.

6. Tan RZH, Lockart I, Abdel Shaheed C, Danta M. Systematic review with meta-analysis: The effects of non-steroidal anti-inflammatory drugs and anti-platelet therapy on the incidence and recurrence of hepatocellular carcinoma. Aliment Pharmacol Ther. 2021;54(4):356–67.

7. Petrick JL, Sahasrabuddhe VV, Chan AT, Alavanja MC, Beane-Freeman LE, Buring JE, et al. NSAID Use and Risk of Hepatocellular Carcinoma and Intrahepatic Cholangiocarcinoma: The Liver Cancer Pooling Project. Cancer Prev Res (Phila). 2015;8(12):1156–62.

8. Pang Q, Jin H, Qu K, Man Z, Wang Y, Yang S, et al. The effects of nonsteroidal anti-inflammatory drugs in the incident and recurrent risk of hepatocellular carcinoma: a meta-analysis. Onco Targets Ther. 2017;10:4645–56.

9. Zappavigna S, Cossu AM, Grimaldi A, Bocchetti M, Ferraro GA, Nicoletti GF, et al. Anti-Inflammatory Drugs as Anticancer Agents. Int J Mol Sci. 2020;21(7).

10. Zhang Z, Chen F, Shang L. Advances in antitumor effects of NSAIDs. Cancer Manag Res. 2018;10:4631–40.

11. Ricciotti E, Wangensteen KJ, FitzGerald GA. Aspirin in Hepatocellular Carcinoma. Cancer Res. 2021;81(14):3751–61.

12. Raza H, John A, Benedict S. Acetylsalicylic acid-induced oxidative stress, cell cycle arrest, apoptosis and mitochondrial dysfunction in human hepatoma HepG2 cells. Eur J Pharmacol. 2011;668(1-2):15–24.

13. Lu L, Sun HC, Zhang W, Chai ZT, Zhu XD, Kong LQ, et al. Aspirin minimized the pro-metastasis effect of sorafenib and improved survival by up-regulating HTATIP2 in hepatocellular carcinoma. PLoS One. 2013;8(5):e65023.

14. Kahraman DC, Bilget Guven E, Aytac PS, Aykut G, Tozkoparan B, Cetin Atalay R. A new triazolothiadiazine derivative inhibits stemness and induces cell death in HCC by oxidative stress dependent JNK pathway activation. Sci Rep. 2022;12(1):15139.

15. Unsal-Beyge S, Tuncbag N. Functional stratification of cancer drugs through integrated network similarity. NPJ Systems Biology and Applications. 2022 Apr 19;8(1):11.

16. Oefinger PE, Bronson DL, Dreesman GR. Induction of hepatitis B surface antigen in human hepatoma-derived cell lines. Journal of General Virology. 1981 Mar;53(1):105–13.

17. Ianevski A, Giri AK, Aittokallio T. SynergyFinder 3.0: an interactive analysis and consensus interpretation of multi-drug synergies across multiple samples. Nucleic Acids Research. 2022 Jul 5;50(W1):W739–43.

18. Subramanian A, Narayan R, Corsello SM, Peck DD, Natoli TE, Lu X, Gould J, Davis JF, Tubelli AA, Asiedu JK, Lahr DL. A next generation connectivity map: L1000 platform and the first 1,000,000 profiles. Cell. 2017 Nov 30;171(6):1437–52.

19. Corsello SM, Bittker JA, Liu Z, Gould J, McCarren P, Hirschman JE, Johnston SE, Vrcic A, Wong B, Khan M, Asiedu J. The Drug Repurposing Hub: a next-generation drug library and information resource. Nature medicine. 2017 Apr;23(4):405–8.

20. Tuncbag N, Gosline SJ, Kedaigle A, Soltis AR, Gitter A, Fraenkel E. Network-based interpretation of diverse high-throughput datasets through the omics integrator software package. PLoS computational biology. 2016 Apr 20;12(4):e1004879.

21. Miao C, Liang C, Tian Y, Xu A, Zhu J, Zhao K, Zhang J, Hua Y, Liu S, Dong H, Zhang C. Overexpression of CAPN2 promotes cell metastasis and proliferation via AKT/mTOR signaling in renal cell carcinoma. Oncotarget. 2017 Nov 11;8(58):97811.

22. Peng X, Yang R, Song J, Wang X, Dong W. Calpain2 upregulation regulates emt-mediated pancreatic cancer metastasis via the Wnt/β-Catenin signaling pathway. Frontiers in medicine. 2022 May 30;9:783592.

23. Nagy Á, Munkácsy G, Győrffy B. Pancancer survival analysis of cancer hallmark genes. Scientific reports. 2021 Mar 15;11(1):6047.

24. Bartha Á, Győrffy B. TNMplot. com: a web tool for the comparison of gene expression in normal, tumor and metastatic tissues. International journal of molecular sciences. 2021 Mar 5;22(5):2622.

25. Yang J, Nie J, Ma X, Wei Y, Peng Y, Wei X. Targeting PI3K in cancer: mechanisms and advances in clinical trials. Molecular cancer. 2019 Dec;18(1):1–28.

26. Durmaz I, Guven EB, Ersahin T, Ozturk M, Calis I, Cetin-Atalay R. Liver cancer cells are sensitive to Lanatoside C induced cell death independent of their PTEN status. Phytomedicine. 2016 Jan 15;23(1):42–51.

27. Kim MS, Kim JE, Lim DY, Huang Z, Chen H, Langfald A, Lubet RA, Grubbs CJ, Dong Z, Bode AM. Naproxen induces cell-cycle arrest and apoptosis in human urinary bladder cancer cell lines and chemically induced cancers by targeting PI3K. Cancer Prevention Research. 2014 Feb 1;7(2):236–45.

28. Xia H, Lee KW, Chen J, Kong SN, Sekar K, Deivasigamani A, Seshachalam VP, Goh BK, Ooi LL, Hui KM. Simultaneous silencing of ACSL4 and induction of GADD45B in hepatocellular carcinoma cells amplifies the synergistic therapeutic effect of aspirin and sorafenib. Cell death discovery. 2017 Sep 11;3(1):1–0.

29. Peng X, Yang R, Song J, Wang X, Dong W. Calpain2 upregulation regulates emt-mediated pancreatic cancer metastasis via the Wnt/β-Catenin signaling pathway. Frontiers in medicine. 2022 May 30;9:783592.

30. Shen C, Yu Y, Li H, Yan G, Liu M, Shen H, Yang P. Global profiling of proteolytically modified proteins in human metastatic hepatocellular carcinoma cell lines reveals CAPN 2 centered network. Proteomics. 2012 Jun;12(12):1917–27.

31. Rouillard AD, Gundersen GW, Fernandez NF, Wang Z, Monteiro CD, McDermott MG, Ma’ayan A. The harmonizome: a collection of processed datasets gathered to serve and mine knowledge about genes and proteins. Database. 2016 Jan 1;2016.

32. Zhang G, Fang T, Chang M, Li J, Hong Q, Bai C, Zhou J. Calpain 2 knockdown promotes cell apoptosis and restores gefitinib sensitivity through epidermal growth factor receptor/protein kinase B/survivin signaling. Oncology Reports. 2018 Oct 1;40(4):1937–46.

33. Ho WC, Pikor L, Gao Y, Elliott BE, Greer PA. Calpain 2 regulates Akt-FoxO-p27Kip1 protein signaling pathway in mammary carcinoma. Journal of Biological Chemistry. 2012 May 1;287(19):15458–65.

34. Li P, Miao C, Liang C, Shao P, Wang Z, Li J. Silencing CAPN2 expression inhibited castration-resistant prostate cancer cells proliferation and invasion via AKT/mTOR signal pathway. BioMed research international. 2017 Feb 9;2017.

