## Supplementary material for "SYNERGISTIC ANTITUMOR EFFECT OF NAPROXEN AND SORAFENIB IN HEPATOCELLULAR CARCINOMA": Table S1, Fig. S1

### SUPPLEMENTARY INFORMATION

**Table S1:** Assessment of synergistic effects (Synergy scores) of all drug combinations involved in this study on HCC cell lines using *Synergyfinder* tool. ND: not determined.

| **Huh7** | Aspirin | Naproxen | Ibuprofen | Flurbiprofen |
|---|---|---|---|---|
| Sorafenib | -16.7 | 16.4 | -4.2 | -6.5 |
| Regorafenib | ND | ND | ND | ND |
| Lenvatinib | ND | ND | ND | ND |

| **Mahlavu** | Aspirin | Naproxen | Ibuprofen | Flurbiprofen |
|---|---|---|---|---|
| Sorafenib | 1.3 | 16.1 | -15.8 | -2.0 |
| Regorafenib | 0.6 | -2.8 | ND | ND |
| Lenvatinib | -14.6 | -11.4 | -11.8 | -7.6 |

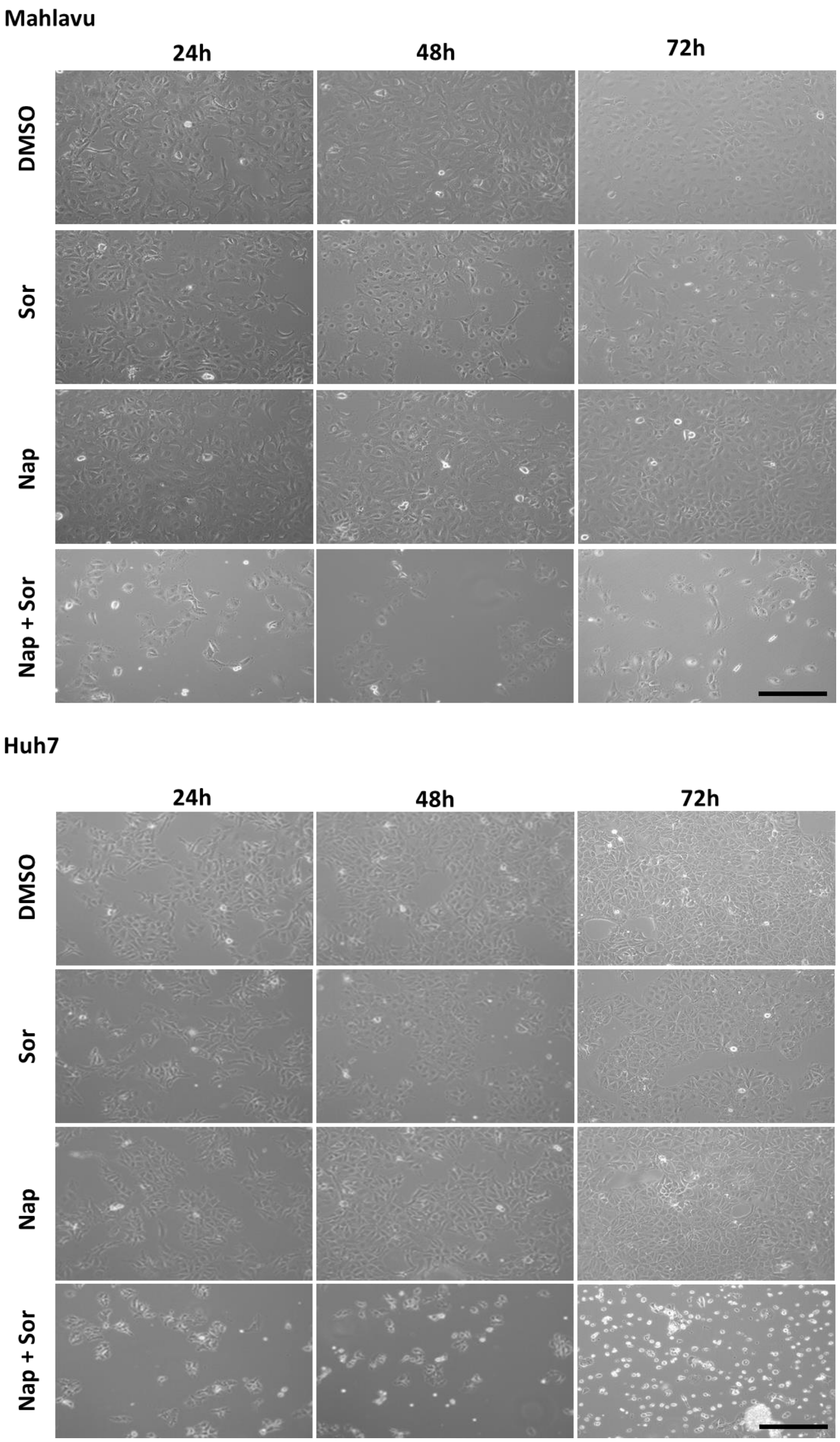


**Figure S1:** Live images of Mahlavu and Huh7 cells after treatment with Sor: 5 μM for Mahlavu, 2.5 μM for Huh7, Nap: 400 μM, or Nap+Sor (5 μM + 400 μM for Mahlavu, and 2.5 μM + 400 μM for Huh7) for 24, 48 or 72 h. Cell images were taken under light microscopy. (Scale bar: 100 μm)
